# Impact of retroactivity on information flows in engineered synthetic biological circuits

**DOI:** 10.64898/2026.03.05.709940

**Authors:** Sailash Moirangthem, Karthik Raman

## Abstract

In biological networks, retroactivity describes the feedback from downstream components that can influence and alter the behavior of upstream systems. This effect poses a major challenge to the modular design of synthetic circuits, where upstream modules are expected to function independently of their connections. Beyond disrupting dynamics, retroactivity can also interfere with how information is transmitted through a network, acting as a bottleneck that reduces the fidelity of signal propagation. Here, we combine stochastic biochemical modeling with information-theoretic analysis to quantify how retroactivity constrains upstream signaling, even in strongly amplified feedback architectures, particularly in the presence of molecular noise. At the same time, we identify parameter regimes in which retroactivity can be exploited as a functional mechanism: downstream loading can trigger controllable state transitions, enabling circuits that respond to changes in their environment or interconnections. These findings suggest design principles for harnessing retroactivity for programmable signal processing and decision-making in cellular computation. Finally, we evaluate feedback-gain tuning as a mitigation strategy and demonstrate that increasing gain alone is insufficient under noisy conditions. We therefore propose complementary approaches to reduce retroactivity and delineate the operating regimes in which each strategy is most effective.

## Introduction

Complex networked dynamical systems are often analyzed by decomposing them into smaller input–output modules, with the assumption that each module preserves its behavior when interconnected. A canonical biological example is the MAPK (Mitogen-Activated Protein Kinase) cascade, where tiers such as Raf, MEK, and ERK can be characterized through well-defined input–output relationships (e.g., phosphorylation states) and then assembled to explain how extracellular cues drive cellular outcomes [1]. This principle of *modularity* is central to both analysis and design of complex systems [2]. In many physical and biological settings, however, modularity breaks down because interconnection alters component dynamics. In genetic networks, for example, connecting a transcriptional regulator to multiple downstream promoters can change its concentration dynamics through binding interactions that “load” the upstream system [3, 4]. This loss of modularity, termed *retroactivity* occurs when downstream components influence upstream dynamics [5]. An analogous loading effect arises in electrical circuits when downstream connections alter upstream voltages or currents. In biomolecular systems (Fig. 1), an activator–repressor clock exhibits sustained oscillations in isolation, but coupling the activator *A* to a downstream system *D* can damp and eliminate oscillations. This breakdown occurs because the binding and unbinding of *A* to promoter sites in the downstream system competes with the upstream biochemical interactions that generate the oscillations. Prior studies quantified retroactivity and proposed “insulation” strategies, typically based on high-gain negative feedback, to attenuate downstream loading and preserve upstream behavior [4–6].

**Figure 1.**
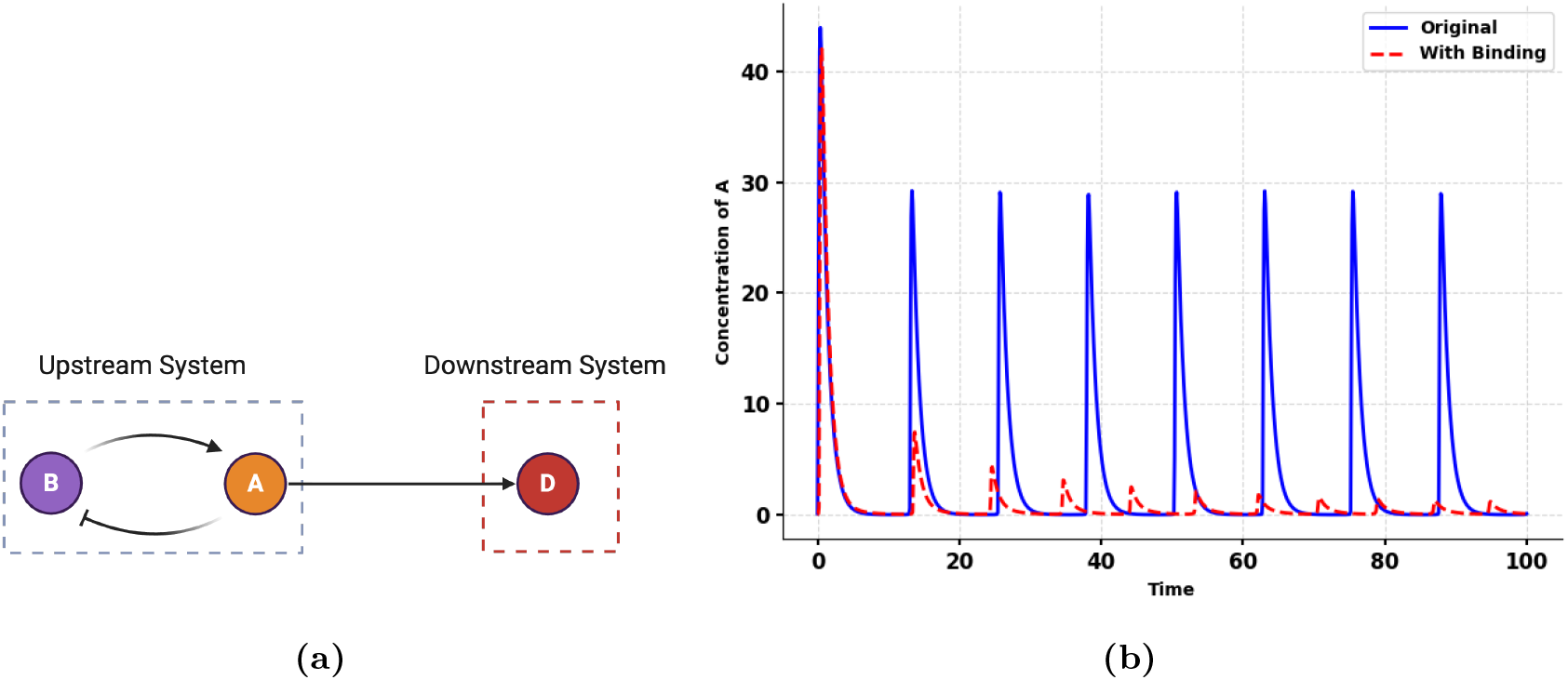
Effect of retroactivity on an oscillatory upstream module. (a) The activator protein *A* forms an oscillatory circuit with *B* (upstream module) and provides timing, driving a downstream component *D*. (b) When the upstream oscillator (“clock”) operates in isolation (solid blue line), A exhibits sustained oscillations. Upon connection to the downstream system (dashed red line), the oscillatory dynamics of *A* are damped and eventually lost.

Despite this progress, important gaps remain. Genetic circuits function through molecular communication [7, 8] : upstream components must reliably influence downstream components not only in their mean behavior, but also in how uncertainty and fluctuations propagate through the network. In this sense, retroactivity is not only a dynamical disturbance as it can degrade the effective communication capacity of a circuit by reducing how predictably downstream species can be inferred or controlled from upstream signals. Yet the impact of retroactivity on directional information flow between molecular species, and how this interacts with intrinsic noise and downstream disturbances, has not been systematically characterized. Moreover, retroactivity is typically treated as an undesirable effect to be suppressed, but downstream loading can in principle be exploited to reshape nonlinear behaviors and induce functional state transitions. Finally, while high-gain negative feedback has been proposed to attenuate retroactivity, whether such insulation strategies preserve information flow under noisy conditions remains unclear.

In this work, we ask: How does downstream loading reshape directional information transfer in biomolecular signaling channels, and when does it effectively act as an information bottleneck? To address this, we develop a unified stochastic and information-theoretic framework to quantify how retroactivity reshapes information transfer in biomolecular networks with single-input/single-output signaling and multiple downstream loads. We measure information flow using directed information [9] and transfer entropy [10]. In the low-copy-number regime, we use Chemical Master Equation (CME) models [11] to capture intrinsic stochasticity; in the high-copy-number regime, we use the linear noise approximation (LNA) [11] to characterize changes in means and covariances. The “low-copy” CME model is exact for discrete molecular counts and captures event-level randomness (reaction jumps, rare transitions). The “high-copy” LNA is obtained as a system-size approximation of the CME and captures small fluctuation propagation (how perturbations/noise are transmitted locally) where fluctuations around the deterministic trajectory become approximately Gaussian and are characterized by a covariance dynamics. Thus, LNA is the asymptotic approximation of CME when copy numbers are large. Across both regimes, we show that retroactivity can substantially reduce effective communication even when state trajectories appear only modestly perturbed, and that feedback-gain tuning alone may fail to prevent information loss in noisy conditions. We further demonstrate that retroactivity can be exploited functionally. For example in toggle-switch circuits, explicit sequestration by downstream binding sites can induce bistability even without cooperative binding, enabling controllable switching through network interconnection [12–14]. We illustrate these results on biologically relevant motifs including toggle switches and activator–repressor oscillators, and discuss implications for modular circuit design, robust cellular computation, and control under molecular noise.

## Results

### Quantifying Information Transfers

#### Low Molecular Counts

To study the effect of retroactivity on the information transfers, we consider an example of a signalling system as shown in Fig. 2 with *N* downstream systems. The system receives an external stimulus as input and, through a cyclic biochemical reaction, transforms it into output signals that regulate various downstream targets or substrates such as DNA binding sites.

**Figure 2.**
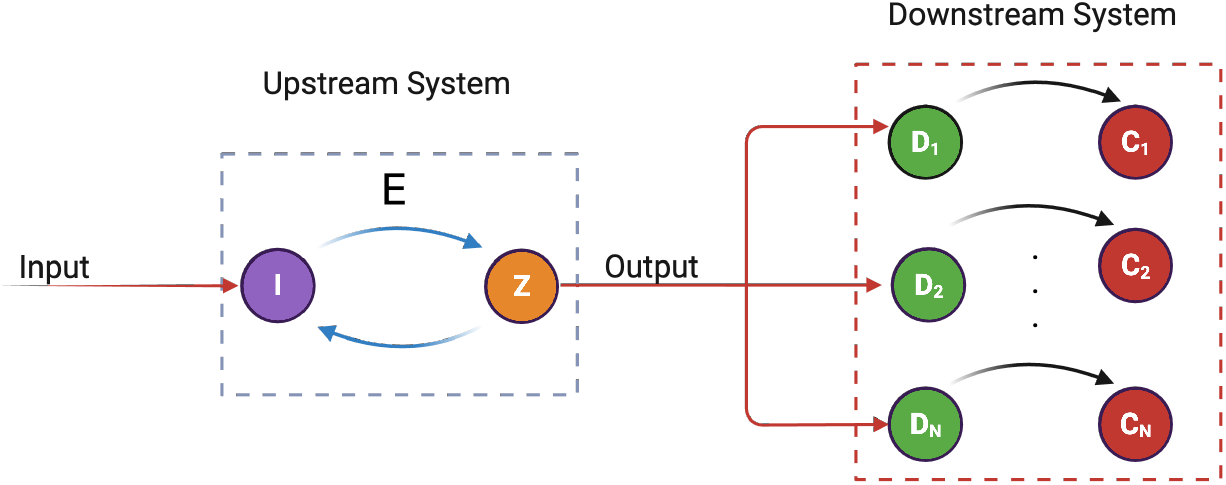
The upstream module (left dotted box) consists of components I and Z with bidirectional interactions (blue arrows indicate mutual regulation). The downstream modules (right dotted box) comprise multiple pairs of components (*D*_1_–*C*_1_ through *D*_*N*_ –*C*_*N*_) receiving inputs from the upstream module (red arrows).

The cyclic biochemical reaction is described by a two-step enzymatic reaction representing the upstream system with no connections to the downstream system as

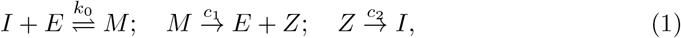

with conservation laws, *E*_tot_ = *E* + *M*, and *I*_tot_ = *I* + *M* + *Z*. The protein *I* binds to an enzyme *E* to form the complex *M* . The complex *M* is transformed into the output protein *Z* at a catalytic rate of *c*_1_. The product *Z* is converted back into the input protein *I* at rate *c*_2_ . The rate constant, 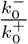, denotes the dissociation constant of the reversible reaction, where 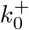 is the forward binding rate constant and 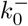 is the backward unbinding of the complex. We refer to the single-input-single-output system model in (1) as a standalone system. We also assume that the upstream system in (1) possesses two distinct symbols, *n*_*I*_ = 2. One symbol is composed of zero molecules, representing the scenario where there is no transmission (*I*(*t*_0_) = 0). While the second symbol is composed of one molecule, indicating the presence of a transmission (*I*(*t*_0_) = 1). Let the initial probability distribution of the input be denoted as 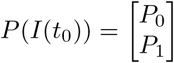, where *P*_1_, *P*_0_ denotes the probability of a transmission and no transmission respectively. When no reaction occurs prior to *t*_0_, *I*_*tot*_ is zero. In this state, there exists only a single microstate, and the system remains in that microstate with a probability of 1. When *I*_*tot*_ = 1 and assuming all reactions occur within a volume Ω, the probability that a reaction will take place within a sufficiently short time interval is determined by propensity functions. The set of chemical reactions can thus be rewritten as a set of transitions between the microstates.

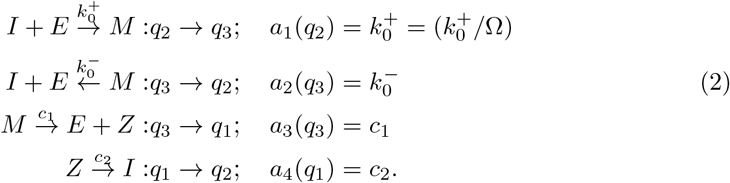

The CME can thus be written using the propensity functions, *a*_*i*_ for each reaction *i* as

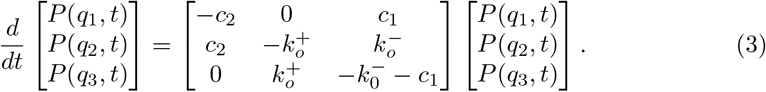

The joint probability mass functions, *P* (*I*^*t*^, *Z*^*t*^) can thus be derived as

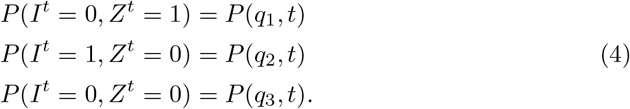

The directed information transfer from the output protein, *Z* to the input protein, *I* is measured using *transfer entropy* [10]. The transfer entropy from *Z* to *I* when the molecular count is low is derived as (Given in Methods 0.1)

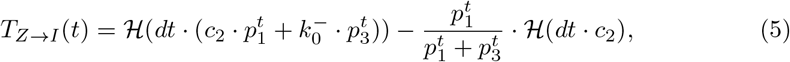

where ℋ (*p*) = −*p* log *p* − (1 − *p*) log(1 − *p*) is the binary entropy function, and *dt* represents an infinitesimal time interval. The term 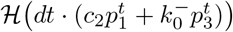 represents the total uncertainty (binary entropy) associated with the possible transition of the input state *I* during an infinitesimal interval *dt*. The probability that *I* changes its state is proportional to the combined rate 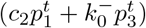, which includes (a) the conversion of *Z* to *I* at rate *c*_2_, weighted by 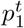, and (b) the unbinding of the complex *M* at rate 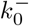, weighted by 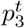 . The term 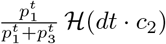 represents the conditional uncertainty in the transitions of *I* that would occur solely due to the conversion of *Z* to *I* with rate *c*_2_, given that the system is in a state where either *I* or *Z* is present (i.e., in the subspace spanned by *q*_1_ and *q*_3_). The weighting factor 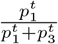 normalizes this uncertainty relative to the combined probability of these states, effectively conditioning on the state of *Z*. Overall, Eq. (5) decomposes the stochastic dynamics of *I* into a mixture of intrinsic reaction uncertainty (due to 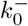) and directed informational influence (due to *c*_2_), providing a quantitative measure of causal feedback in the biochemical cycle.

Given there is no cyclic conversion, i.e., *c*_2_ = 0 from *Z* to *I*, then *T*_*Z*→*I*_ (*t* → ∞) → 0. When *c*_2_ = 0, the only transition of *I* (from 0 to 1) is due to 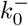 and is given by *q*_3_ ↔ *q*_2_. The cyclic transitions *q*_3_ ↔ *q*_2_ do not contribute to the transfer entropy *T*_*Z*→*I*_ because they occur entirely within the subspace where *Z*^*t*^ = 0. Therefore, 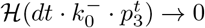 as *t* → ∞ (*i. e*.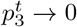) at steady state when there is no activation path (*i*.*e*., *c*_2_ = 0). Also, if 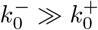, then there is a more complex dissociation to *I* resulting in an increased influence of *Z* on *I* through the intermediate complex *M* . Thus, an increase in 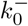 leads to an increase in *T*_*Z*→*I*_ (*t*). In the steady state at *t*_*s*_, *T*_*Z*→*I*_ (*t*) is given by (also see Methods)

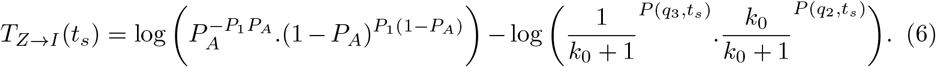

Note that when *c*_2_ = 0, we have *P*_*A*_ = 1, *P* (*q*_3_, *t*_*s*_) = 0 and *P* (*q*_2_, *t*_*s*_) = 0 and therefore *T*_*Z*→*I*_ (*t*_*s*_) = 0. Also, when 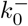 is much faster than 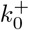, the second term in (6) goes to zero as *P* (*q*_3_, *t*_*s*_) ≈ 0 and *k*_0_ ≫ 1. Under this condition, 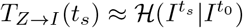 and reaches its maximum value.

#### High Molecular Counts

To study the effect of retroactivity on the transfer entropy between species when there is a higher number of molecules, we consider a transcriptional model with a one-step enzymatic reaction as studied in [15] and shown in Fig. 3.

**Figure 3.**
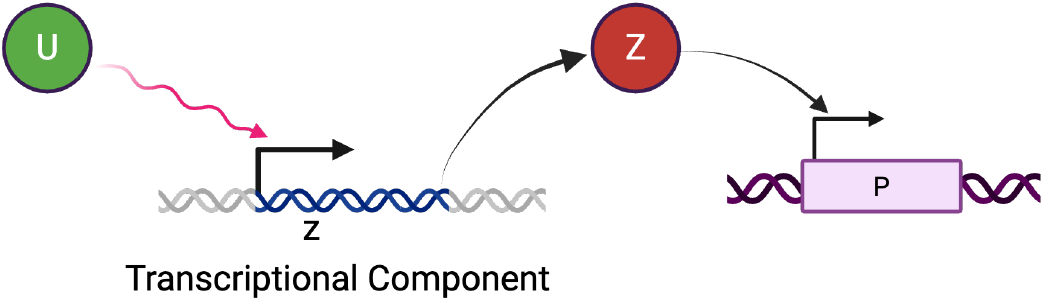
Transcriptional component with *U* as input and producing *Z* as output. A downstream component takes *Z* as the input through its reversible binding to the promoter *p*.

The activity of the promoter controlling gene *z* depends on the amount of *U* bound to the promoter. The biochemical process can be written as

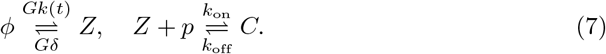

The time varying function *k*(*t*) describes the transcription rate of a gene *z* which expresses protein *Z* and *δ* defines the degradation rate of protein *Z*. The amplification gain *G* is assumed to be tunable to attenuate the retroactivity due to the downstream system. We also assume that *k*(*t*) and *δ* are of the same order and that the production and the degradation processes are slower than the binding and the unbinding reactions, that is *k*_off_ ≫ *Gδ* and *k*_off_ ≫ *Gk*(*t*) [16]. Denoting *p*_*T*_ as the total concentration of the promoter *p*, the differential equation model of the process in (7) can be written as

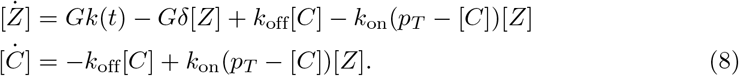

Comparing the dynamics of the isolated and the connected systems, the dynamics of *Z* is affected by an additional rate of change *s* = *k*_off_[*C*] − *k*_on_(*p*_*T*_ − [*C*])[*Z*]. Thus, when *s* = 0, we get the isolated system dynamics. Previous work [4] shows how the retroactivity effect can be compensated by increasing the gain *G*. To study the effect of retroactivity on the communication pattern between molecules, we consider the stochastic model for (8).

Let *P* (*Z, C, p*; *t*) denote the probability that the number of molecules of species Z, C, and p are *Z, C* and *p*, respectively, at time *t*, given that the initial probabilities at *t* = *t*_0_ are *Z*_0_, *p*_0_ and *C*_0_, respectively. If Ω denotes the volume of the system, then the number of molecules of the species are given by *Z* = Ω[*Z*] and *C* = Ω[*C*]. Using Ω-expansion [11], we expand *Z* as

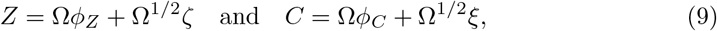

where *ζ, ξ* are the stochastic components and *ϕ*_*Z*_, *ϕ*_*C*_ are the deterministic components. The resulting equations for *ϕ*_*Z*_, *ϕ*_*C*_ and *ζ, ξ* are found using the linear noise approximation. The evolution of *ϕ*_*Z*_, *ϕ*_*C*_ are given by

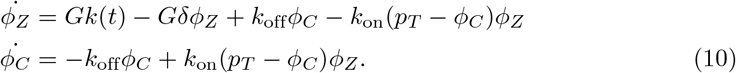

The information transfer from the downstream system *C* to the upstream *Z* is computed from the evolution of the stochastic components *ζ, ξ* and is given by (also see Methods)

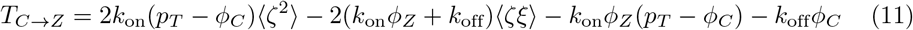

The measure *T*_*C*→*Z*_ can also be understood as the reduction in the uncertainty of the concentration *Z* due to the binding to the promoter site *p*. When there is no binding to the promoter site, *k*_off_ = *k*_on_ = 0 and therefore ⟨*ζ*^2^⟩_no binding_ = ⟨*ζ*^2^⟩_binding_ and from (11), *T*_*C*→*Z*_ = 0. Another important property of *T*_*C*→*Z*_ is the asymmetrical information transfer, that is, *T*_*C*→*Z*_≠ *T*_*Z*→*C*_.

### Effects of retroactivity on information transfers

#### Retroactivity in Low Molecular Counts

Each downstream system in Fig. 2 is composed of a reversible reaction

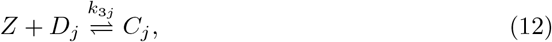

with conservation law 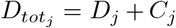. Here the output protein of the upstream system in (1) binds with *D*_*j*_ to form the complex *C*_*j*_. The downstream system (12) perturbs these dynamics by sequestering the output species *Z* into complexes *C*_*j*_. This modifies the conservation law of the upstream network as

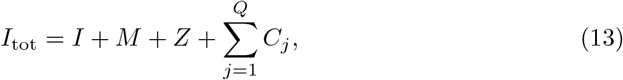

where *Q* denotes the number of downstream modules coupled to the upstream system. The additional term ∑_*j*_ *C*_*j*_ effectively reduces the amount of free *Z* available, thereby altering the transition probabilities 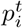 and the corresponding information transfer between species. In the presence of downstream systems, it can be shown that the term *c*_1_ approximates the impact of retroactivity due to the downstream system and for *N* downstream systems, 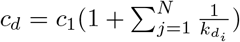, where 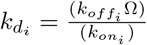 is the dissociation constant of *i*^*th*^ downstream, with 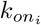 and 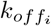 denoting the forward binding and the backward unbinding rates. Note that when there is no downstream system, *c*_*d*_ = *c*_1_. Fig. 4a shows the difference in the evolution of probabilities for the standalone system and the system with a downstream channel.

**Figure 4.**
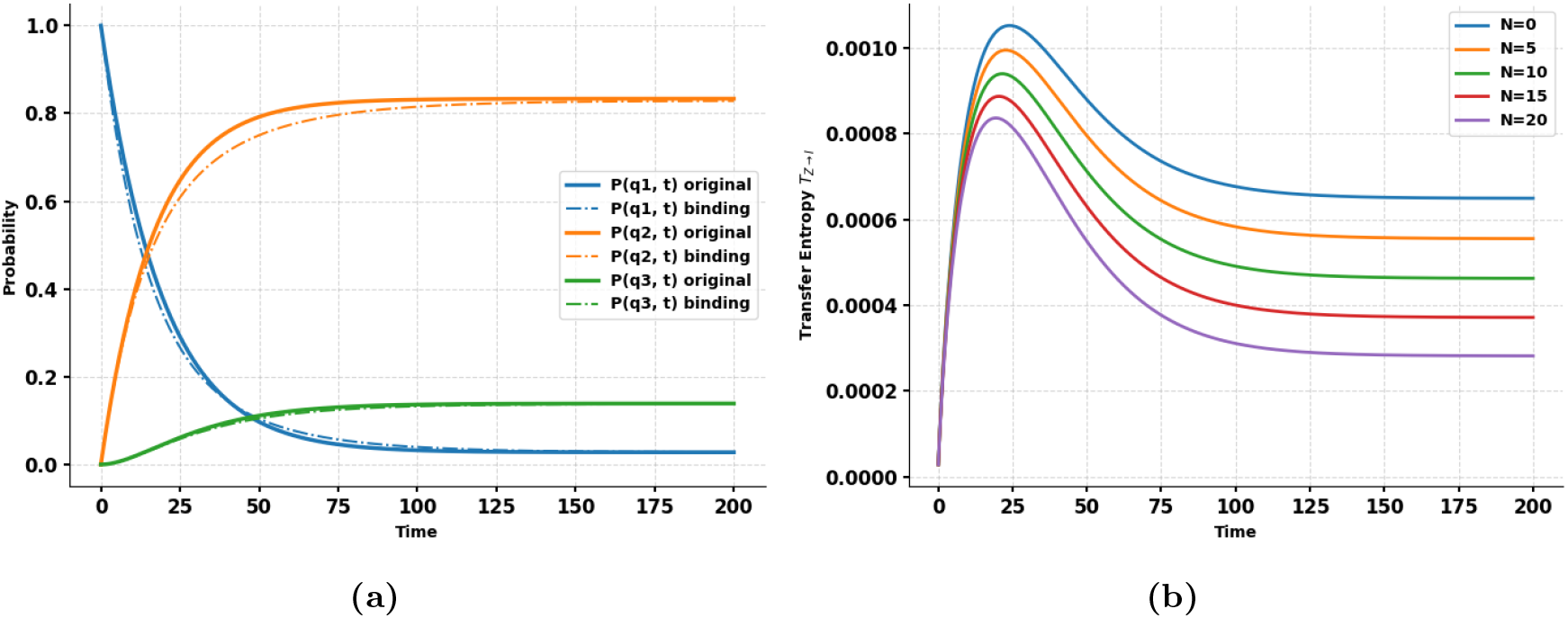
(a) Effect of retroactivity on the evolution of probabilities. The probabilities *P* (*q*_*i*_, *t*) denote the probabilities for the standalone system, and *P*_*r*_(*q*_*i*_, *t*) denotes the probabilities for the system with a downstream channel. (b) Effect of retroactivity on *T*_*Z*→*I*_ for multiple downstream systems. For the simulation, we take *c*_1_ = 0.01(*nM/s*), *c*2 = 0.05*s*^−1^, 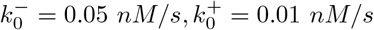 and 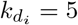. With values adopted from [8].

The transfer entropy curves for an increasing number of downstream systems are shown in Fig. 4b. In all plots, the transfer entropy *T*_*Z*→*I*_ starts at a low value, rises to a peak, and then gradually decreases to a steady-state value 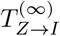. In the beginning, *Z* is synthesized from *M* and starts influencing *I*. Early on, the history of *Z* contains significant predictive information about the future state of *I*, so transfer entropy rises. The peak occurs when *Z* has its strongest causal effect on *I*, i.e., when knowledge of *Z* gives maximal predictive power over *I*. As the system approaches steady state, the dynamics become less dependent on the initial conditions and interactions become more “memoryless.” At this stage, *I*’s future is largely determined by its current state rather than the history of *Z*, so the additional information from *Z* declines, causing TE to decrease. Adding more downstream systems effectively “dilutes” the influence of *Z* on *I*. When *N* is very large, almost all molecules of *Z* interact with downstream systems instead of *I*, so *Z* loses its predictive power over *I*, and *T*_*Z*→*I*_ approaches zero. We can draw a similar conclusion from the fact that, for infinite downstream systems, the probability of the number of free molecules of *Z* being transformed to protein *I* tends to zero and therefore H(*I*^*t*+1^|*I*^*t*^, *Z*^*t*^) → H(*I*^*t*+1^|*I*^*t*^) as *N* → ∞. Similarly, in the steady state, the effect of retroactivity due to downstream systems is captured by *P*_*A*_ and *P*_*B*_ (see Methods, (31) and (33)). In the presence of *N* downstream systems, we replace *c*_1_ by *c*_*d*_ and the modified probabilities are

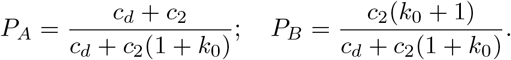

As the number of downstream systems increases, *c*_*d*_ → ∞ and *P*_*A*_ → 1, *P*_*B*_ → 0, and using (6), *T*_*Z*→*I*_ (*t*_*s*_) → 0.

#### Retroactivity in High Molecular Counts

In the deterministic case, the terms *k*_off_[*C*] − *k*_on_(*p*_*T*_ − [*C*])[*Z*] in (8) represent the retroactivity to the output and captures how the binding dynamics to the downstream promoter affects the upstream concentration *Z* and can be seen as a mass (or signal) flow from the upstream module to the downstream module due to binding/unbinding dynamics. It acts as a feedback load on the upstream system. The second equation models the input stage of the downstream system. The effect of the retroactivity on the behavior of *Z* can be significant and is illustrated in Fig. 5a. Fig. 5b shows how the retroactivity can be attenuated by increasing the gain *G*.

**Figure 5.**
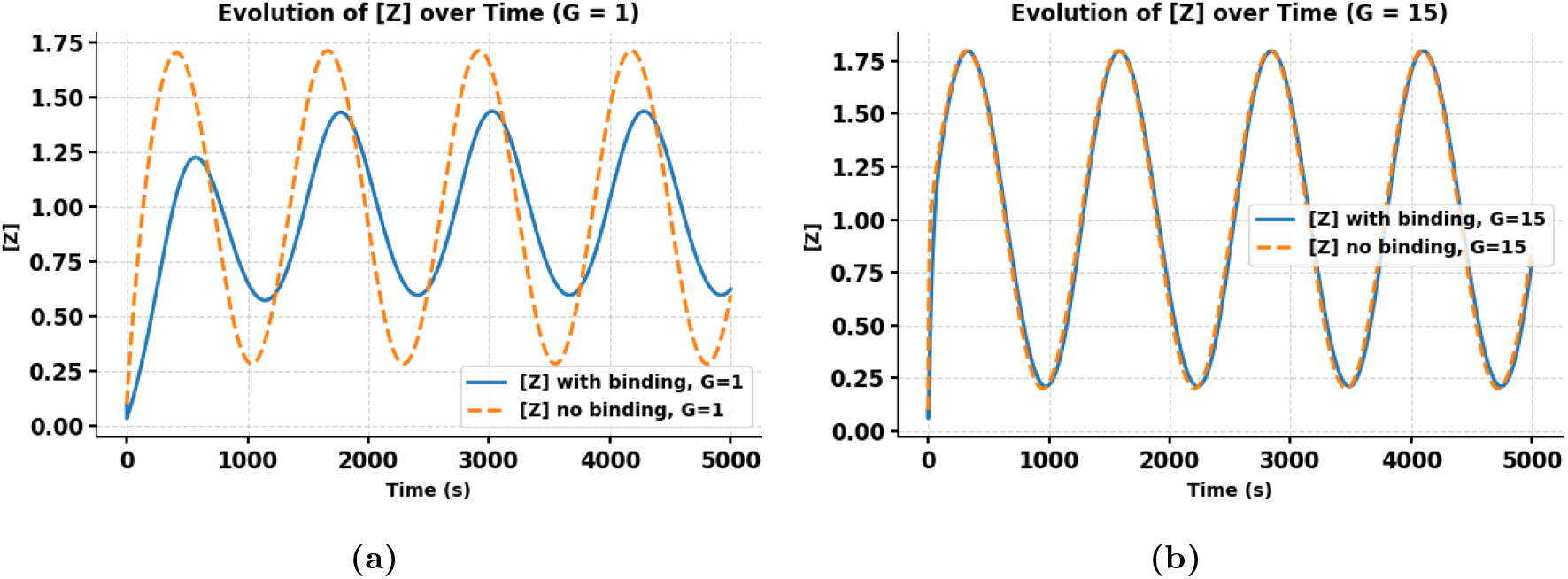
Time evolution of the concentration of species *Z* under different scaling factors and binding conditions. Here, we choose *ω* = 0.005,*δ* = 0.01*s*^−1^, *k*_*d*_ = 20M, *k*_off_ = 50*s*^−1^, and *k*(*t*) = *δ*(1 + 0.8 sin *ωt*)M*s*^−1^ with values adopted from [15]. (a) For *G* = 1, *Z* dynamics are shown both with reversible binding to downstream targets (solid line) and without binding (*k*_on_ = *k*_off_ = 0, dashed line). Binding moderates the oscillatory response and reduces the peak amplitude. (b) For *G* = 15, the evolution of *Z* is similarly shown with and without binding. Increasing the scaling factor amplifies the oscillations of *Z* and compensates the effect due to downstream binding. These plots illustrate how downstream interactions and system scaling influence the temporal behavior of *Z*.

In the deterministic framework [5], increasing the gain *G* enhances the steady-state level of the signaling species *Z*, thereby compensating for the reduction in [*Z*] caused by the downstream load. However, in the stochastic regime, the same strategy becomes insufficient because the intrinsic noise terms scale with *G*. Specifically, in the linear noise approximation, the variance of *Z* is proportional to the effective production and degradation fluxes, ⟨*ζ*^2^⟩ ∼ *G*(*k*(*t*) + *δ*[*Z*]), implying that amplification of the mean signal by increasing *G* also amplifies its stochastic fluctuations. As a result, the signal-to-noise ratio 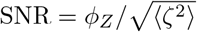 saturates or decreases with large *G* [17–19]., limiting the effective information transfer between *Z* and downstream molecules. Although the damping coefficient in the variance dynamics (Eq. 39) appears to increase with G via the term (*Gδ* + *k*_on_(*p*_*T*_ − *ϕ*_*C*_)), the retroactivity-induced feedback depends nonlinearly on *G*. As *G* increases, the mean level *ϕ*_*Z*_ also rises, enhancing sequestration and reducing the fraction of unbound binding sites (*p*_*T*_ − *ϕ*_*C*_). This weakens the retroactivity-associated feedback term *k*_on_(*p*_*T*_ − *ϕ*_*C*_), while the noise-injection terms *Gk*(*t*) and *Gδϕ*_*Z*_ continue to grow, further amplifying fluctuations in *Z*. Thus, beyond a certain point, increasing *G* alone cannot suppress retroactivity because the effective damping against noise decreases despite higher gain.

### Attenuating retroactivity

In low molecular counts, the inverse of the dissociation constant 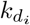 measures the *affinity* of *Z* binding to downstream systems. Therefore, a high 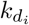 represents a *low affinity* between *Z* and *i*^*th*^ downstream site. Thus, as 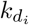 increases, the effect of retroactivity decreases and *c*_*d*_ → *c*_1_, the catalytic rate of the standalone system. Therefore, an approach to attenuate the effect of retroactivity is to increase the dissociation constants 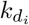 to all the downstream systems. Fig. 7 shows compensation for the decrease in *T*_*Z*→*I*_ due to the retroactivity effect by increasing 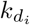.

In the high molecular counts, if both the input amplification (*G*) and the degradation rate (*δ*) are sufficiently large, the contribution of the retroactivity-related term −2*k*_on_(*p*_*T*_ − *ϕ*_*C*_)⟨*ζ*^2^⟩ in *d*⟨*ζ*^2^⟩*/dt* ((39) in Methods) becomes negligible compared to the dominant damping term −2*Gδ*⟨*ζ*^2^⟩. Moreover, the cross-correlation and offset terms 2(*k*_on_*ϕ*_*Z*_ + *k*_off_) ⟨*ζξ*⟩, *k*_on_*ϕ*_*Z*_(*p*_*T*_ − *ϕ*_*C*_), and *k*_off_*ϕ*_*C*_ are largely independent of *G* and *δ*. Hence, as *G* and *Gδ* increase, the dynamics of the standalone and connected systems converge, and from Eq.(11), the information transfer *T*_*C*→*Z*_ → 0. This behavior, shown in Fig.6b, illustrates how enhancing both amplification and degradation attenuates the effects of retroactivity and restores modular signaling behavior.

**Figure 6.**
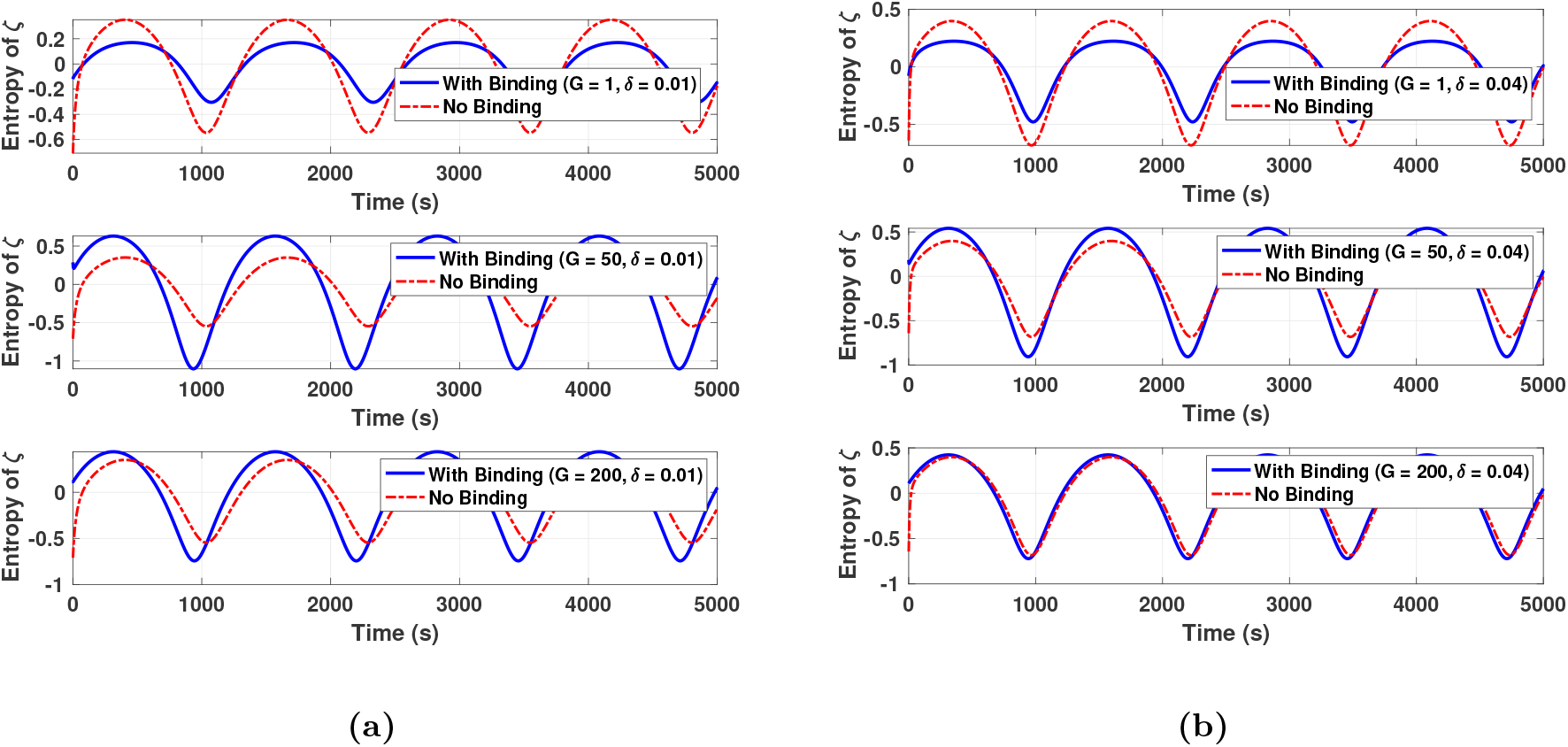
(a) Effect of retroactivity due to the downstream system. For these plots, we use *ω* = 0.005,*δ* = 0.01*s*^−1^, *k*_*d*_ = 20M, *k*_off_ = 50*s*^−1^, and *k*(*t*) = *δ*(1 + 0.8 sin *ωt*)M*s*^−1^. (b) Compensating for the effect of retroactivity by amplifying the input signal and enhancing the degradation of *δ* = 0.05.

**Figure 7.**
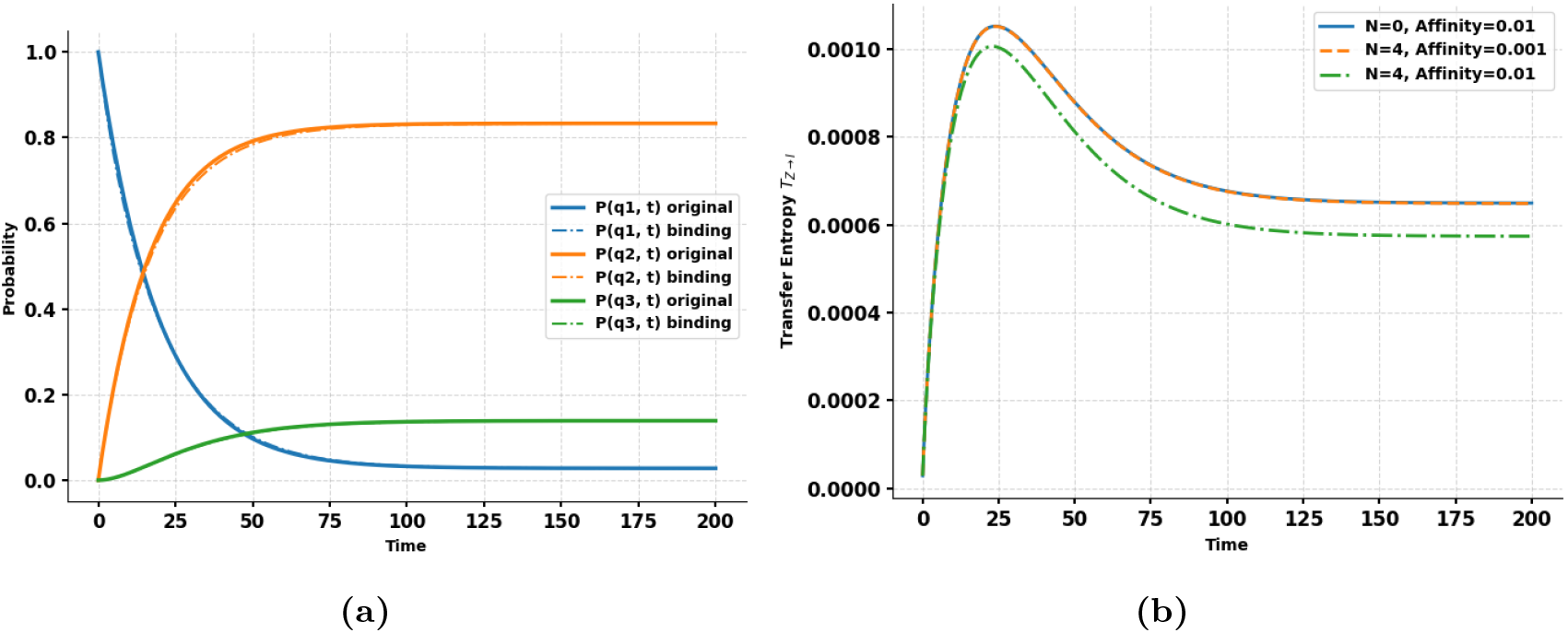
(a) Effect of retroactivity on the evolution of probabilities. The probabilities *P* (*q*_*i*_, *t*) denote the probabilities for the standalone system, and *P*_*r*_(*q*_*i*_, *t*) denotes the probabilities for the system with a downstream channel. (b) Effect of retroactivity on the transfer entropy from *Z* to *I* decreases as the affinity 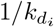 decreases from 0.2 to 0.04 for *N* = 4 downstream systems connected to *Z*.

### Retroactivity in a biological toggle switch

A toggle switch is a basic genetic circuit composed of two proteins, A and B, with concentrations denoted by *A* and *B*, respectively [12, 20]. Each protein represses the synthesis of the other: protein B inhibits the production of protein A by binding in *n* copies to the promoter of gene A, and similarly, protein A represses the production of protein B by binding to its promoter. The system exhibits two stable equilibrium states. Once a cell settles into one of these states, it remains there despite small perturbations. Consequently, the toggle switch serves as a canonical model for studying cellular decision-making and fate determination [21–25].

We begin with the canonical reduced toggle-switch model in (14), which captures mutual repression using effective Hill-type production terms and does not explicitly represent promoter binding/unbinding reactions. We use the standard Michaelis–Menten/Hill-type toggle-switch model [26] (Eq. 14)

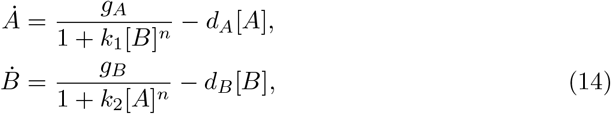

where *k*_1_, *k*_2_ represent the repression strengths, *n* is the Hill coefficient, *g*_*A*_ and *g*_*B*_ are the maximal production rates, and *d*_*A*_ and *d*_*B*_ are the degradation rates of proteins A and B, respectively. In this reduced description, the toggle is modular only in isolation: its input–output behavior is defined assuming that the transcription factors regulating each promoter are not significantly sequestered by other loads. Retroactivity arises when the toggle is embedded in a larger circuit and one of its transcription factors is connected to downstream binding sites. In that case, a fraction of the transcription factor becomes bound downstream, so the repression in (14) must depend on the free transcription factor. This interconnection creates a feedback from the downstream binding state to the upstream toggle dynamics, which is the defining feature of retroactivity. The switch can be controlled by varying the repression strengths. In *E. coli* cells, the proteins A and B are representations of LacI and TetR, fused to the fluorescent reporters mKate2 (Red Fluorescent Protein, RFP) and monomeric Enhanced Green Fluorescent Protein, mEGFP (GFP), respectively. The repression strengths *k*_1_ and *k*_2_ can be controlled dynamically by addition of the diffusible molecules, isopropyl-*β*-D-thiogalactopyranoside (IPTG) and anhydrotetracycline (aTc), respectively. The cells switch to an RFP-dominant state (or GFP-dominant state) upon addition of aTc (or IPTG, respectively) and retains the state after washout of the inducer as shown in Fig. 8.

**Figure 8.**
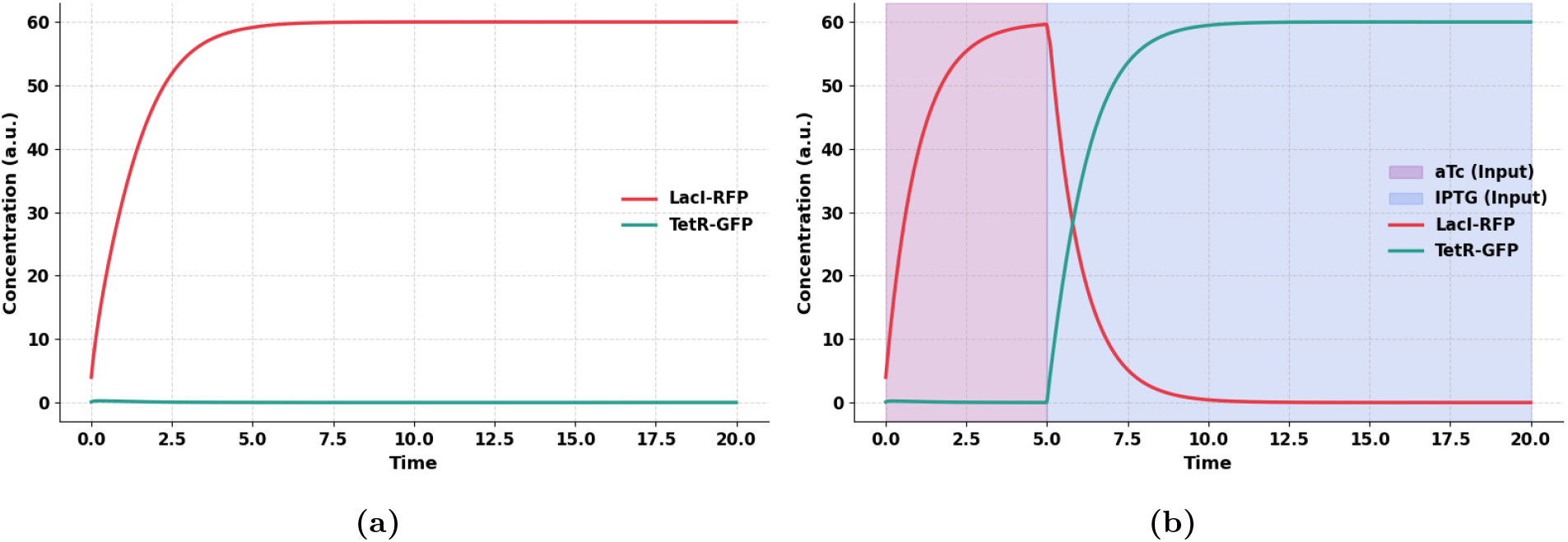
Simulated toggle switch model in (14), with *g*_*A*_ = *g*_*B*_ = 50, *k*_1_ = *k*_2_ = 1, *n* = 2, *d*_*A*_ = *d*_*B*_ = 1 b) Uncontrolled toggle switch with *k*_1_, *k*_2_ as constants, i.e, *k*_1_ (aTc) = 1, and *k*_2_(IPTG) = 1 (a) Controlling the toggle switch by varying *k*_1_ (aTc) and *k*_2_ (IPTG). We use *k*_1_ = 0.1 for *t* ∈ [0, 5] and 1 for *t* ∈ [5, 20] and *k*_2_ = 0.1 for *t* ∈ [5, 20] and 1 for *t* ∈ [0, 5]. More details are given in Supplementary Note 2.

Time-evolving transfer entropies between LacI and TetR when the system is externally controlled using aTc and IPTG pulses are shown in Fig. 9a. Derivation of transfer entropy for the toggle switch is given in Supplementary Note 1. The transfer entropy is nearly symmetric in both directions as the system is in a transient phase with both genes actively repressing each other. Fig. 9b shows the time evolution of delayed transfer entropy in both directions for uncontrolled systems (no pulses), with different TetR production rates (*g*_*B*_). When repression strengths (or production rates) differ significantly, the resulting dynamical asymmetry increases fluctuations and sensitivity in the system. This enhanced activity leads to higher symmetric transfer entropy in both directions. In contrast, when *g*_*A*_ ≈ *g*_*B*_, the system exhibits more balanced and less variable dynamics, resulting in lower symmetric transfer entropy.

**Figure 9.**
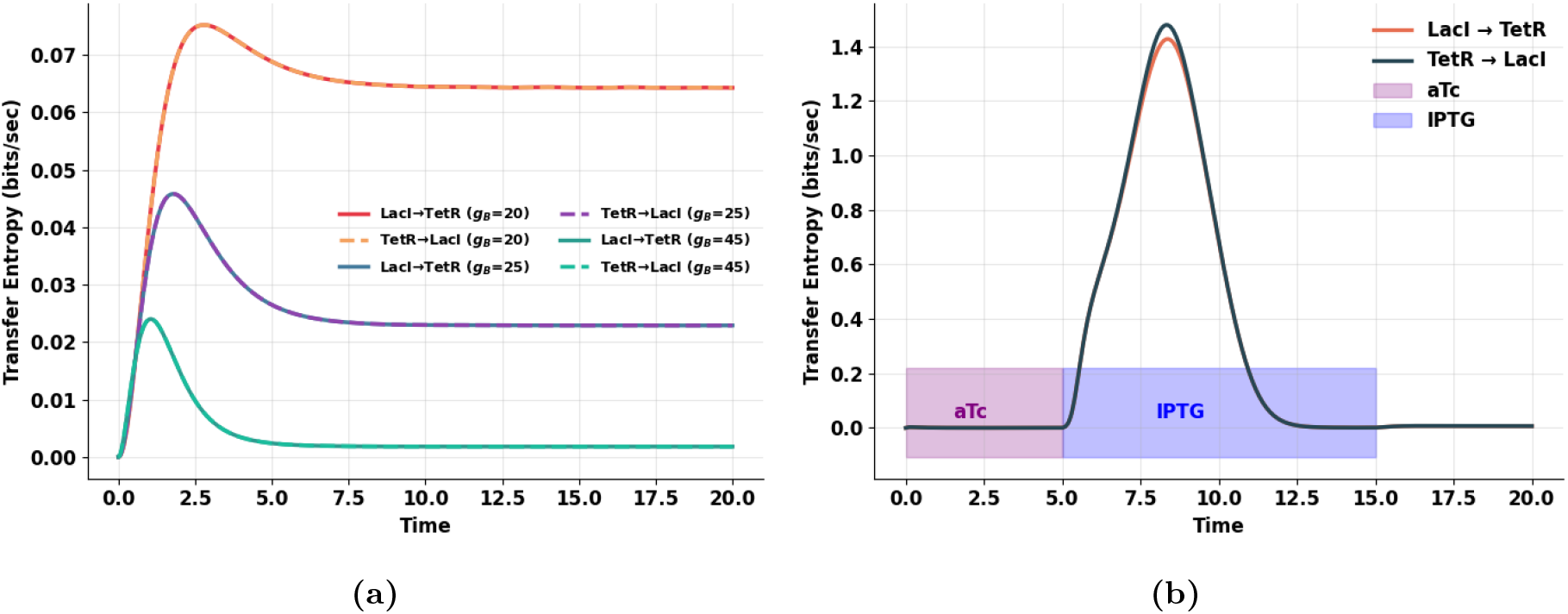
Simulated toggle switch model in (14), with *g*_*A*_ = 50, *k*_1_ = *k*_2_ = 1, *n* = 2, *d*_*A*_ = *d*_*B*_ = 1, *β* = 2, *η* = 2, *γ* = 1. (a) Transfer entropy in an uncontrolled toggle switch (b) Transfer Entropy in the controlled Switch. The purple region (t ∈ [0, 5]) shows when the aTc pulse is applied, thereby affecting TetR repression. The blue region (t ∈ [5, 15]) shows when the IPTG pulse is applied, thereby repressing the LacI concentration.

### Retroactivity-Modified Toggle Switch Model

Although the toggle switch model we study is a reduced description without explicit binding reactions, it is treated as an upstream module whose output drives a downstream single-input single-output channel. Retroactivity therefore enters not within the internal reactions of the toggle itself, but through this interconnection. The downstream system sequesters a fraction of the upstream transcription factor, effectively altering the free concentration that participates in the toggle’s regulatory interactions. Even in reduced models, such loading can shift nullclines, modify switching thresholds, and change the effective bistability landscape. Thus the toggle remains modular when isolated, but loses modularity when connected to downstream targets because of this load-induced feedback, which is the defining feature of retroactivity.

To analyze the effect of retroactivity on information transfer, we extended the canonical toggle switch model in (14) by explicitly incorporating the interaction of the upstream transcription factor *B* with downstream modules. When *B* is connected to a downstream module, a fraction of it becomes bound to the downstream binding sites (or promoter regions), creating a complex *C*. Denoting the total concentration of downstream sites as [*S*]_*T*_, the reversible binding reaction 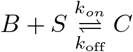 introduces two new fluxes corresponding to association and dissociation. The total concentration of *B* is now partitioned as 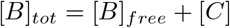, where only the free form participates in repression of *A*.

Accordingly, the dynamics are modified as

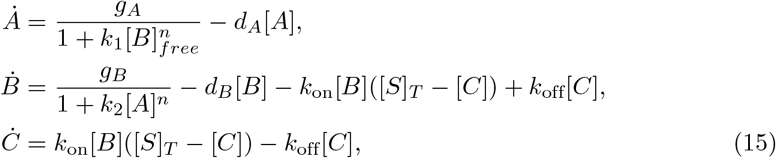

with [*B*]_*free*_ = [*B*] − [*C*]. This formulation captures the *retroactive load* imposed by the downstream connection. The binding–unbinding terms in Eq. (15) introduce a feedback from the downstream system to the upstream dynamics, effectively reducing the concentration of free *B* available for regulation. As a result, the apparent repression strength of *B* on *A* weakens, and the system exhibits slower response and reduced bistability compared to the isolated toggle switch. In the stochastic setting, these additional terms are also reflected in the Linear Noise Approximation (LNA) through the augmented Jacobian and diffusion matrices, allowing the resulting covariance evolution to quantify how retroactivity modulates the transfer entropy *TE*_*B*→*A*_ (see Methods section). Fig. 10 illustrates the impact of retroactivity on both the dynamics of a genetic toggle switch (Fig. 10a) and the corresponding transfer entropy (TE) in both directions (Fig. 10b). Before *t* = 10, TetR concentration (blue) rises rapidly to its steady state, while LacI (red) remains repressed near zero due to strong TetR-mediated inhibition. At *t* = 10, retroactivity is introduced by coupling the TetR output to 20 downstream binding sites. This acts as an additional load on TetR, effectively reducing the free concentration of TetR. There’s a transient dip in TetR concentration due to binding to downstream sites, but the system quickly stabilizes and returns to a new steady state. LacI remains repressed due to the high TetR level, so the toggle remains in the same expression state. In Fig. 9b, before *t* = 10, there is a peak in transfer entropy during the transient phase (early dynamics), indicating strong information flow as the system settles into its stable state. The transfer entropy values are symmetric because the toggle dynamics are reciprocal (mutual inhibition), and the LNA model assumes Gaussian noise. After *t* = 10, the introduction of retroactivity slightly increases the transfer entropy at the steady state. However, because the expression state does not switch, the transfer entropy remains symmetric and relatively low.

**Figure 10.**
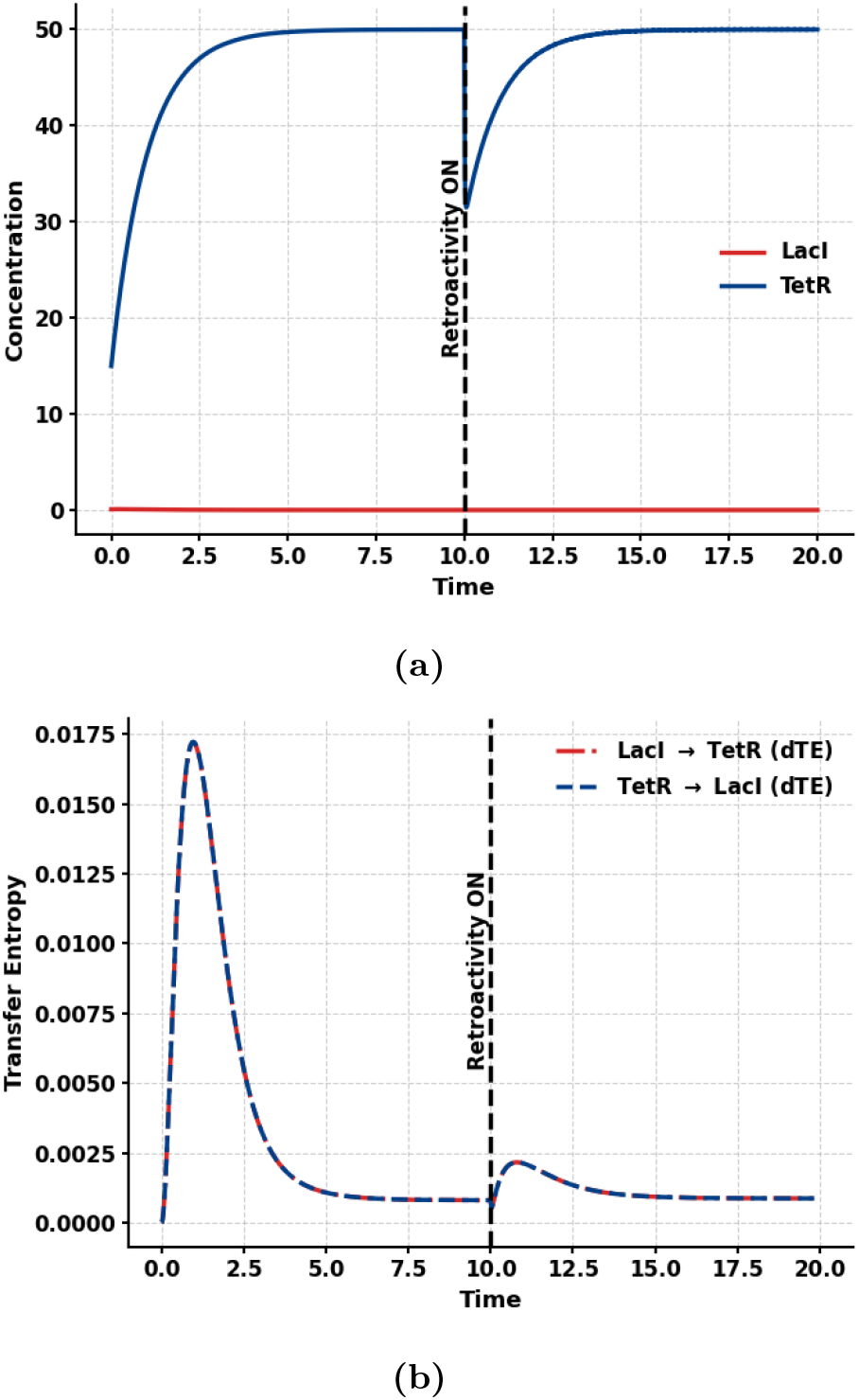
Simulated toggle switch model with *g*_*A*_ = *g*_*B*_ = 50, *β* = 2, *η* = 2, *γ* = 1, *k*_1_ = *k*_2_ = 1, *k*_on_ = 2, *k*_off_ = 1 (a) Concentration changes due to 20 downstream systems (b) Effect of retroactivity on Transfer Entropy in both directions

Fig. 11 shows that introducing retroactivity by coupling TetR to 70 downstream targets at *t* = 10 induces a switch in the toggle circuit from TetR dominance (blue) to LacI dominance (red). This switch occurs because retroactivity reduces the free TetR available to repress LacI. The corresponding transfer entropy (right panel) sharply peaks during the transition, reflecting a burst of causal influence as the system reorganizes. After switching, transfer entropy stabilizes at a new steady state. This highlights how retroactivity can go beyond being a problem to contend with in synthetic circuits, but can be leveraged to drive state transitions and modulate information flow in genetic circuits.

**Figure 11.**
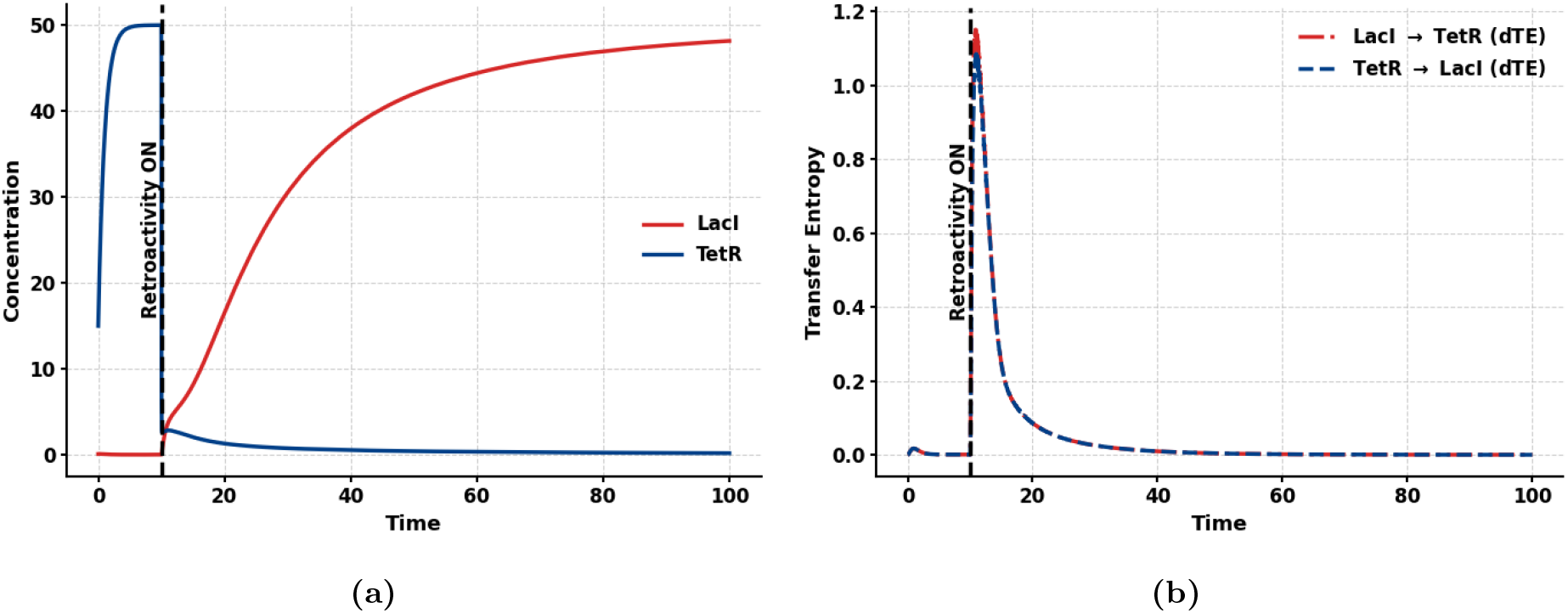
Effect of Retroactivity on Toggle Switch Dynamics and Information Flow. The values of the parameters are *g*_*A*_ = 50, *g*_*B*_ = 50, *β* = 2, *η* = 2, *γ* = 1, *k* = 1, *k*_off_ = 1, *k*_on_ = 2. (a) Time evolution of LacI and TetR concentrations in a synthetic toggle switch. At *t* = 10, retroactivity is introduced by coupling TetR to 70 downstream binding sites, reducing its free concentration and triggering a switch to LacI dominance. (b) Transfer entropy (dTE) in both directions increases sharply during the switching event, indicating a transient rise in causal influence between the genes. After the transition, TE stabilizes at a new steady-state level.

#### Retroactivity for detecting disturbance

Fig. 12 highlights how a transient disturbance affects the toggle switch dynamics and its associated information transfer. While the concentrations of LacI and TetR (left) show minimal deviation upon the onset of disturbance at *t* = 10, the differential transfer entropy (right) exhibits a clear and immediate response. This reveals that the causal influence between the two genes becomes temporarily altered, even though their steady-state behavior remains largely unchanged. The information-theoretic measure thus provides a more sensitive and quantitative view of the system’s internal adaptation, capturing subtle dynamic changes that are not visible in the state trajectories alone.

**Figure 12.**
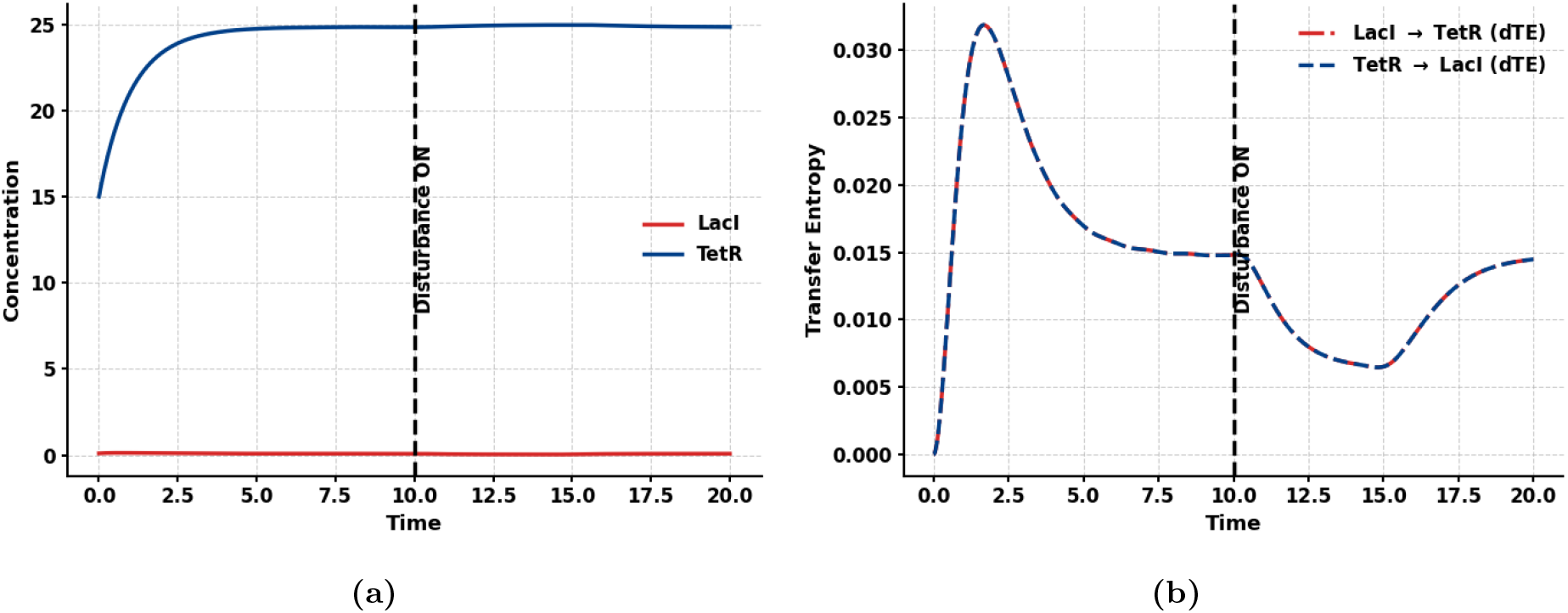
Effect of Disturbance on Toggle Switch Dynamics and Information Flow. (a) Time evolution of LacI and TetR protein concentrations in a toggle switch circuit with parameters *g*_*A*_ = 50, *g*_*B*_ = 25, *β* = 2, *η* = 2, *γ* = 1, *k* = 1. A disturbance is applied at *t* = 10 that changes *g*_*B*_ = 24. TetR shows a slight transient deviation, while LacI remains suppressed. The impact of the disturbance is not visually prominent in the concentrations. (b) A distinct dip in information transfer follows the disturbance onset at *t* = 10, indicating a transient loss of directional predictability. This highlights the higher sensitivity of information-theoretic measures in detecting system perturbations.

#### Retroactivity as a toggle switch controller

Figures 13 illustrate how retroactivity can serve as a dynamic control input for modulating toggle switch behavior. By turning retroactivity ON and OFF at specific time intervals, the system’s state and the flow of information between components can be selectively influenced. These results suggest that retroactivity, when appropriately timed, can act as a control signal.

**Figure 13.**
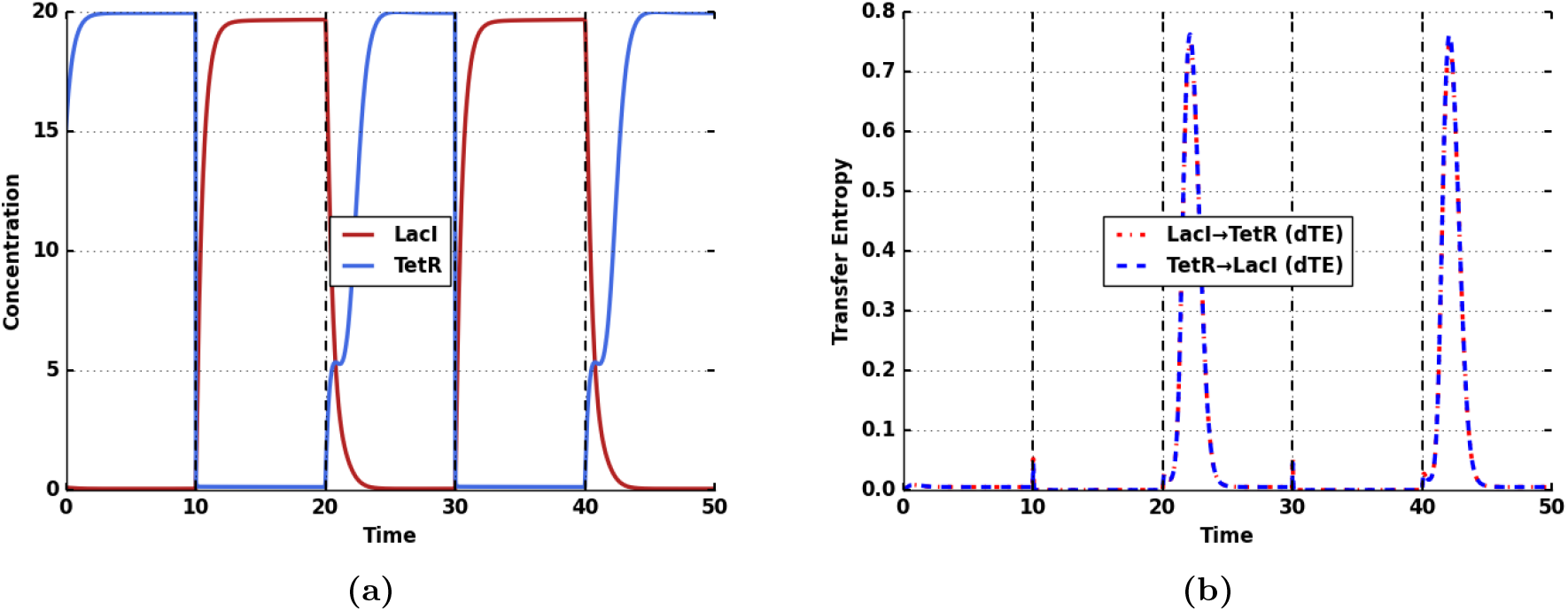
(a) Dynamic response and directional information transfer in a toggle switch with periodically modulated retroactivity. Time evolution of LacI (red) and TetR (blue) concentrations under switching retroactivity applied during *t* ∈ [10, 20] and *t* ∈ [30, 40] (vertical dashed lines). The parameter values are *g*_*A*_ = 40, *g*_*B*_ = 40, *β* = 2, *η* = 2, *γ* = 2, *k* = 1. (b) The red dash-dotted curve (LacI → TetR) and blue dashed curve (TetR → LacI) represent the instantaneous directional information transfer computed from the Linear Noise Approximation. Peaks in differential transfer entropy (dTE) appear immediately after each switching event at *t* = 10, 20, 30, and 40 s,reflecting transient bursts of directional influence as the system switches under the new retroactivity condition. The small dips following each peak indicate brief losses of predictive coupling during these transitions.

## Discussion

This work establishes an information-theoretic framework for understanding how retroactivity alters communication between molecular components in biochemical networks. Rather than focusing solely on concentration dynamics, we quantify how retroactive coupling modifies the direction and magnitude of information flow between interacting molecules. This perspective provides a fundamental shift in how retroactivity is analyzed, moving from state-based to information-based descriptions that directly capture the causal influence one molecule exerts on another. By employing transfer entropy and directed information, we quantify how the coupling of molecular modules affects their ability to transmit and process information. These measures extend traditional deterministic and stochastic analyses by revealing hidden dependencies and causal asymmetries that emerge when upstream and downstream systems are interconnected. Unlike correlation-based or gain-based metrics, TE and directed information remain valid in nonlinear, noisy regimes, providing a general framework that unifies stochastic and deterministic models under a single quantitative measure of influence.

In the low-copy-number regime, where intrinsic noise dominates, we model the system through the chemical master equation and compute transfer entropy directly from the evolving probability distributions. This approach captures how retroactivity disrupts the fidelity of signal propagation at the molecular level. The reduction in transfer entropy observed with increasing retroactive load indicates a loss of effective communication between upstream regulators and their targets, a phenomenon consistent with the loss of modularity described by Del Vecchio et al. [5] but now revealed through an information-theoretic lens. Importantly, transient peaks in transfer entropy during retroactivity induced switching reveal temporary reorganizations of control within the network, changes that are not visible from concentration dynamics alone.

In the high-copy-number regime, our use of the Fokker–Planck equation and covariance-based transfer entropy links retroactivity to changes in how uncertainty and fluctuations spread through the network. This continuous approximation shows that retroactivity reshapes the network’s variability and alters its information structure. By relating covariance dynamics to directional information transfer, our framework connects retroactivity analysis with principles from system theory and stochastic thermodynamics, complementing earlier studies that describe retroactivity as a dynamical load [6, 27].

The information-theoretic viewpoint also suggests new design principles. In the stochastic regime, weakening binding affinity can restore information transmission by minimizing retroactive feedback on probability fluxes. For the high molecular count in the deterministic framework, increasing the gain compensate the reduction in the concentration of the upstream molecule caused by the downstream load [3, 15]. In our study, we show that increasing the gain alone cannot suppress retroactivity because of the noise term that scales with the gain.

Overall, this study reframes retroactivity not merely as a dynamical perturbation but as an information bottleneck that constrains communication between biomolecular subsystems. Transfer entropy and directed information provide powerful diagnostic tools for revealing when and how such bottlenecks form, evolve, or can be exploited for functional purposes.

Beyond the specific context of synthetic gene circuits, the information-theoretic framework developed here has broad implications for analyzing communication and control in complex biological systems. The ability to quantify directional information transfer directly from molecular interactions provides a scalable approach for uncovering hidden regulatory dependencies in larger genetic, signaling, or metabolic networks. Similar to how transfer entropy has illuminated causal relations in neuronal and ecological systems, applying it to biomolecular circuits can reveal how information is routed, buffered, or dissipated in response to environmental or evolutionary pressures. This cross-disciplinary perspective bridges molecular systems biology and network information theory, offering a powerful conceptual and computational toolset for studying how living systems maintain coordinated yet modular information processing across scales, from genes to cells to tissues.

## Conclusion

In this work, we developed an information-theoretic approach to study how retroactivity affects the behavior of synthetic genetic circuits. By analyzing both the dynamics and the transfer entropy between components, we showed that retroactivity can alter the direction and strength of information flow, often leading to switching or instability in toggle switches. Our results highlight that transfer entropy provides a more sensitive indicator of these effects than concentration profiles alone, especially in stochastic regimes. We also found that increasing the production gain of upstream components can help mitigate retroactivity, though this strategy has limits in noisy systems. The methods presented here are most relevant for transcriptional and post-translational networks where modules interact through shared species or binding sites.

These ideas can be extended to more complex systems, including spatially distributed networks, circuits with feedback-insulated modules, or those with adaptive gain control. Importantly, our results suggest that retroactivity itself can be used as a design feature in cellular computation. By tuning the number and nature of downstream connections, we show that the behavior of a toggle switch—such as whether or not it switches states—can be externally modulated. This opens up possibilities for building programmable responses into synthetic circuits by leveraging retroactive effects. Ultimately, this work lays the groundwork for designing robust, context-aware communication in multicellular and distributed synthetic systems.

## Methods

### Computing Information Transfers in Low Molecular Regime

To compute the information transfer using (16) when the molecular count is low, we first solve the Chemical Master Equation (CME), which governs the probabilities of microstates in a chemical reaction system. A microstate is defined as the number of molecules of each species in a stochastic reaction system. Assuming no reaction starts before an initial time, *t*_0_, we denote *I*_*tot*_(*t*_0_) = *I*(*t*_0_). We assume that the upstream system in (1) possesses two distinct symbols, *n*_*I*_ = 2. One symbol is composed of zero molecules, representing the scenario where there is no transmission (*I*(*t*_0_) = 0). While the second symbol is composed of one molecule, indicating the presence of a transmission (*I*(*t*_0_) = 1). Let the initial probability distribution of the input be denoted as 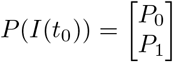, where *P*_1_, *P*_0_ denote the probability of a transmission and no transmission, respectively. The directed information transfer from the output protein, *Z*, to the input protein, *I*, is measured using *transfer entropy* [10]. Transfer entropy from *Z* to *I* is defined as the reduction in the uncertainty of future values of *I* given past values of *I* and *Z*. That is, transfer entropy from *Z* to *I* is given by

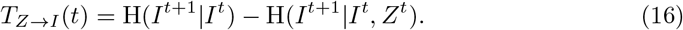

To evaluate *T*_*Z*→*I*_, we need to find *P* (*I*^*t*+1^|*I*^*t*^) and *P* (*I*^*t*+1^|*I*^*t*^, *Z*^*t*^), where *P* (*I*^*t*+1^|*I*^*t*^) is the probability that the particle is in state *I* at time *t* + 1 given that it was in state *I* at time *t*. And *P* (*I*^*t*+1^|*I*^*t*^, *Z*^*t*^) is the probability that the particle is in state *I* at time *t* + 1 given that it was in state (*I, Z*) at time *t*.

When no reaction occurs prior to *t*_0_, *I*_*tot*_ is zero. In this state, only a single microstate exists, and the system remains in that microstate with a probability of 1. When *I*_*tot*_ = 1 and assuming all reactions occur within a volume Ω, the probability that a reaction will take place within a sufficiently short time interval is determined by propensity functions. The set of chemical reactions can thus be rewritten as a set of transitions between the microstates.

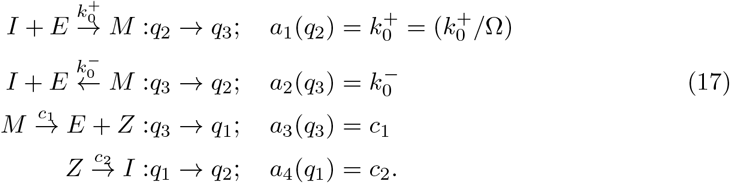

The CME can thus be written using the propensity functions, *a*_*i*_ for each reaction *i*, as

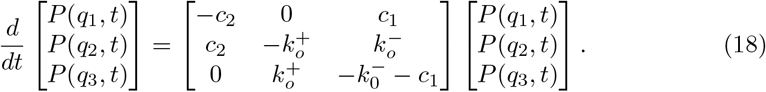

The joint probability mass functions, *P* (*I*^*t*^, *Z*^*t*^) can thus be derived as

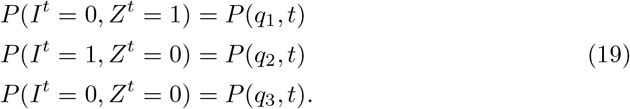

To compute, *H*(*I*^*t*+1^ = 1|*I*^*t*^), we discretize (18) so that

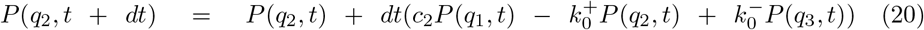

Equation (20) can be decomposed as sum of *P* (*I*^*t*+*dt*^ = 1 | *I*^*t*^ = 0) and *P* (*I*^*t*+*dt*^ = 1 | *I*^*t*^ = 1), where

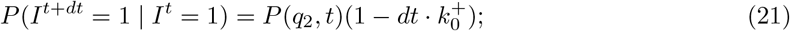

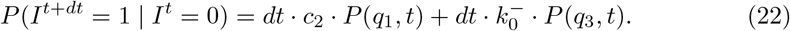

The term 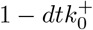 represents the probability that no transition occurs within the small time interval *dt*. We define three different cases of information transfer for different values of *P* (*I*^*t*^) and *P* (*Z*^*t*^).

#### Case I

*I*^*t*^ = 1: The particle is in state *I* at time *t* and corresponds to the microstate *q*_2_ from (17) and (19). From the chemical reaction model, transitions away from *I*^*t*^ = 1 (i.e., from *q*_2_ to *q*_3_) happen via 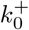, and this transition does not depend on *Z*^*t*^. Since the future of *I* depends only on the current value *I*^*t*^ = 1, and not on *Z*^*t*^, *P* (*I*^*t*+*dt*^ |*I*^*t*^ = 1, *Z*^*t*^) = *P* (*I*^*t*+*dt*^ |*I*^*t*^ = 1) and using (16), *T*_*Z*→*I*_ (*t*) = 0.

#### Case II

*I*^*t*^ = 0, *Z*^*t*^ = 1: In this case, the system is at microstate *q*_1_ and the only possible transition is 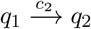. This transition probability depends directly on *Z*^*t*^ through the state *q*_1_, where the transition rate *c*_2_ is active only when *Z*^*t*^ = 1. Therefore, using (22), we can write

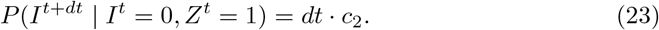

#### Case III

*I*^*t*^ = 0, *Z*^*t*^ = 0: In this case, the system is at microstate *q*_3_ and the only possible transition is 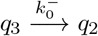. This transition probability depends directly on *Z*^*t*^ through the state *q*_3_, where the transition rate 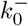 is active only when *Z* Therefore, using (22), we can write

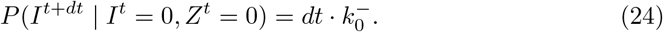

Using (16), transfer entropy from *Z*^*t*^ to *I*^*t*+*dt*^ for Case II is

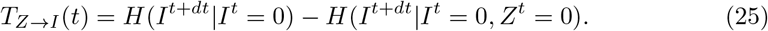

From the definition (22), the expression for the conditional probability of the particle in state *I*^*t*+*dt*^, given *I*^*t*^ = 0 and *Z*^*t*^ = 1, is:

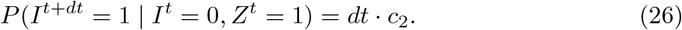

Using (25) and (26), we can write

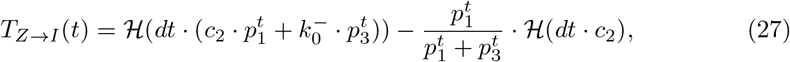

where ℋ (*p*) = −*p* log *p* − (1 − *p*) log 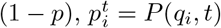.

The derivation of the transfer entropy for Case III (*I*^*t*^ = 0, *Z*^*t*^ = 0) follows the same structure as in Equation (27), with the second term replaced by 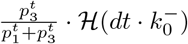, reflecting the contribution from transitions occurring solely due to *Z*^*t*^ = 0.

#### Steady State transfer entropy

The steady state solutions of (18) are given by

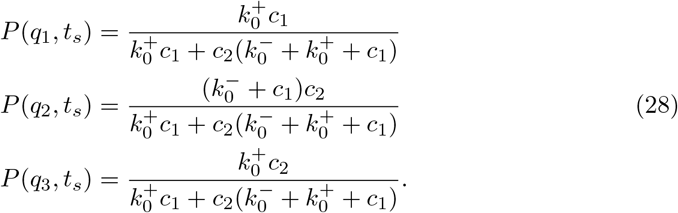

The transfer entropy from *t*_0_ to *t*_*s*_ is given by

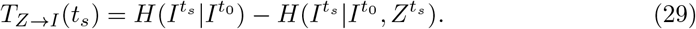

To compute (29), we first derive 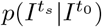 from (28) by marginalizing over 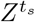 from the joint distribution, 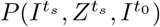. This results in the following transition matrix:

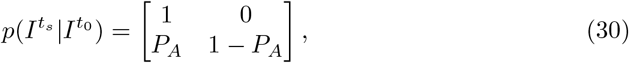

where each row corresponds to the initial state 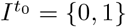 and each column to the resulting state 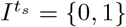. Here, 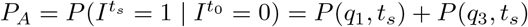, represents the total probability of transitioning to *I* = 1 at time *t*_*s*_, whether *Z* = 1 or *Z* = 0. At steady state, 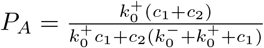. Denote 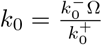, where Ω is the volume in which the reactions take place, and assume 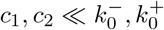, so that

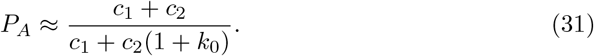

Similarly, the joint *pmf*, 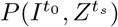 can be found by marginalizing over 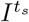 and using *P* (*I*(*t*_0_)) as

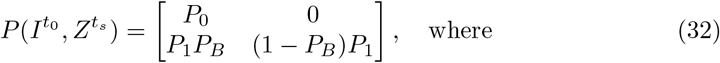

*P*_*B*_ = *P* (*q*_2_, *t*_*s*_) + *P* (*q*_3_, *t*_*s*_) and can be approximated as

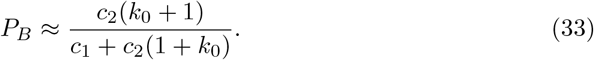

Using (30), we have the expression,

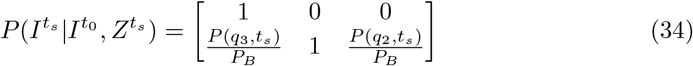

where the first row corresponds to the initial state 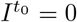 and the second row is when 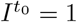. The columns correspond to the values of 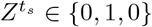, and their impact on 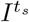. The first term in the second row is the probability 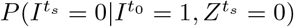. Using (29), the transfer entropy at the steady state is written as

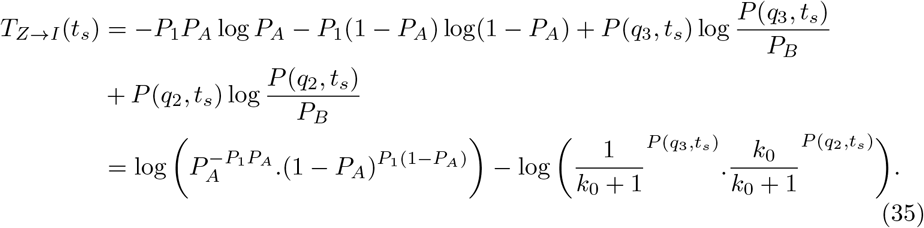

### Computing Information Transfers in High Molecular Regime

Let *P* (*Z, C, p*; *t*) denote the probability that the number of molecules of species Z, C, and p are *Z, C* and *p*, respectively, at time *t*, given that the initial probabilities at *t* = *t*_0_ are *Z*_0_, *p*_0_ and *C*_0_, respectively. If Ω denotes the volume of the system, then the number of molecules of the species is given by *Z* = Ω[*Z*] and *C* = Ω[*C*]. Using Ω-expansion [11], we expand *Z* as

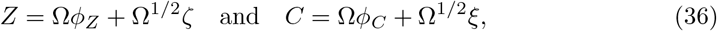

where *ζ, ξ* are the stochastic components and *ϕ*_*Z*_, *ϕ*_*C*_ are the deterministic components. The resulting equations for *ϕ*_*Z*_, *ϕ*_*C*_ and *ζ, ξ* are found using the linear noise approximation. The evolution of *ϕ*_*Z*_, *ϕ*_*C*_ are given by

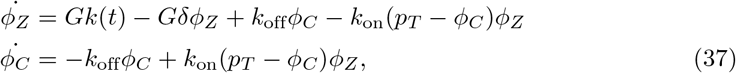

and the resulting Fokker-Planck equation is given by

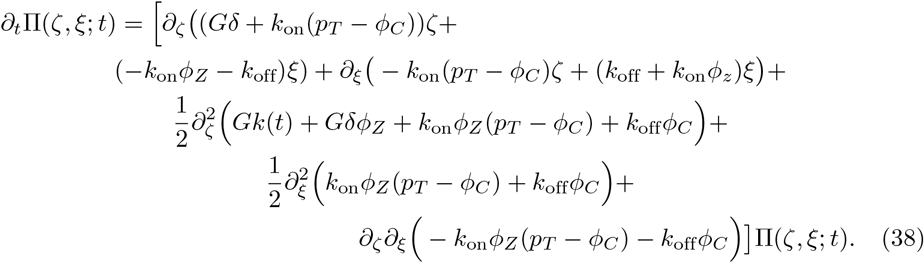

The Fokker–Planck equation can generally be written as:

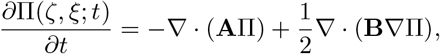

where **A** is the drift vector and **B** is the diffusion tensor. The drift terms for *ζ, ξ*, are identified in the first-order derivatives with respect to *ζ, ξ* and the diffusion coefficients are identified from the second-order derivative terms. Using the drift and the diffusion terms, the equations of second moments are given as

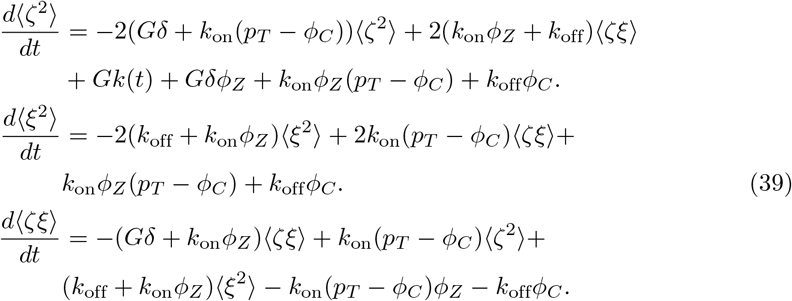

To quantify the information transfer from the downstream system *C* to the upstream *Z*, we examine the change in entropy of the stochastic component of *Z*, in the presence and absence of the downstream component *C*. We define this time-varying quantity as the *information transfer* in the sense of Liang–Kleeman’s information transfer [28] and is defined as

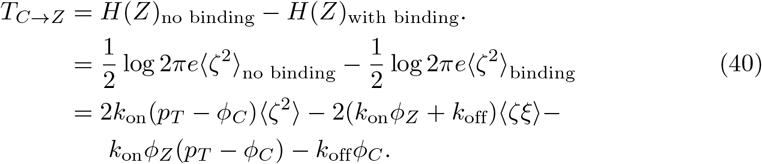

The measure *T*_*C*→*Z*_ can also be understood as the reduction in the uncertainty of the concentration *Z* due to the binding to the promoter site *p*. When there is no binding to the promoter site, *k*_off_ = *k*_on_ = 0 and therefore ⟨*ζ*^2^⟩_no binding_ = ⟨*ζ*^2^⟩_binding_ and from (40), *I*_*C*→*Z*_ = 0. Another important property of *I*_*C*→*Z*_ is the asymmetrical information transfer, that is, *I*_*C*→*Z*_≠ *I*_*Z*→*C*_.

